# Fine-scale human population structure in southern Africa reflects ecogeographic boundaries

**DOI:** 10.1101/098095

**Authors:** Caitlin Uren, Minju Kim, Alicia R. Martin, Dean Bobo, Christopher R. Gignoux, Paul D. van Helden, Marlo Möller, Eileen G. Hoal, Brenna Henn

**Affiliations:** SA MRC Centre for TB Research, DST/NRF Centre of Excellence for Biomedical Tuberculosis Research, Division of Molecular Biology and Human Genetics, Faculty of Medicine and Health Sciences, Stellenbosch University, Cape Town, 8000; Department of Ecology and Evolution, Stony Brook University, Stony Brook, NY 11794; Analytic and Translational Genetics Unit, Department of Medicine, Massachusetts General Hospital, Boston, MA 02114; Program in Medical and Population Genetics, Broad Institute of Harvard and MIT, Cambridge, MA 02142; Department of Genetics, Stanford University, Stanford, CA 94305

**Keywords:** Ancestry, Population Structure, KhoeSan, Pastoralism

## Abstract

Recent genetic studies have established that the KhoeSan populations of southern Africa are distinct from all other African populations and have remained largely isolated during human prehistory until about 2,000 years ago. Dozens of different KhoeSan groups exist, belonging to three different language families, but very little is known about their population history. We examine new genome-wide polymorphism data and whole mitochondrial genomes for more than one hundred South Africans from the ≠Khomani San and Nama populations of the Northern Cape, analyzed in conjunction with 19 additional southern African populations. Our analyses reveal fine-scale population structure in and around the Kalahari Desert. Surprisingly, this structure does not always correspond to linguistic or subsistence categories as previously suggested, but rather reflects the role of geographic barriers and the ecology of the greater Kalahari Basin. Regardless of subsistence strategy, the indigenous Khoe-speaking Nama pastoralists and the N|u-speaking ≠Khomani (formerly hunter-gatherers) share ancestry with other Khoe-speaking forager populations that form a rim around the Kalahari Desert. We reconstruct earlier migration patterns and estimate that the southern Kalahari populations were among the last to experience gene flow from Bantu-speakers, approximately 14 generations ago. We conclude that local adoption of pastoralism, at least by the Nama, appears to have been primarily a cultural process with limited genetic impact from eastern Africa.

**Data deposition:** Data files are freely available on the Henn Lab website: http://ecoevo.stonybrook.edu/hennlab/data-software/

**Summary:** Distinct, spatially organized ancestries demonstrate fine-scale population structure in southern Africa, implying a more complex history of the KhoeSan than previously thought. Southern KhoeSan ancestry in the Nama and ≠Khomani is shared in a rim around the Kalahari Desert. We hypothesize that there was recent migration of pastoralists from East Africa into southern Africa, independent of the Bantu-expansion, but the spread of pastoralism within southern Africa occurred largely by cultural diffusion.

## Introduction

The indigenous populations of southern Africa, referred to by the compound ethnicity “KhoeSan” (Schlebusch 2010), have received intense scientific interest. This interest is due both to the practice of hunter-gatherer subsistence among many groups – historically and to present-day – and genetic evidence suggesting that the ancestors of the KhoeSan diverged early on from all other African populations (Behar *et al.* 2008; Tishkoff *et al.* 2009; Henn *et al.* 2011, 2012; Pickrell *et al.* 2012; Veeramah *et al.* 2012; Barbieri *et al.* 2013). Genetic data from KhoeSan groups has been extremely limited until very recently, and the primary focus has been on reconstructing early population divergence. Demographic events during the Holocene and the ancestry of the Khoekhoe-speaking pastoralists have received limited, mostly descriptive, attention in human evolutionary genetics. However, inference of past population history depends strongly on understanding recent population events and cultural transitions.

The KhoeSan comprise a widely distributed set of populations throughout southern Africa speaking, at least historically, languages from one of three different linguistic families – all of which contain click consonants rarely found elsewhere. New genetic data indicates that there is deep population divergence even among KhoeSan groups (Pickrell *et al.* 2012; Schlebusch *et al.* 2012, 2013; Schlebusch and Soodyall 2012; Barbieri *et al.* 2013), with populations living in the northern Kalahari estimated to have split from southern groups 30,000-35,000 years ago (Pickrell *et al.* 2012; Schlebusch *et al.* 2012; Schlebusch and Soodyall 2012). Pickrell et al. (2012) estimate a time of divergence between the northwestern Kalahari and southeastern Kalahari population dating back to 30,000 years ago; “northwestern” refers to Juu-speaking groups like the !Xun and Ju/’hoansi while “southeastern” refers to Taa-speakers. In parallel, Schlebusch et al. (2012) also estimated an ancient time of divergence among the KhoeSan (dating back to 35,000 ya), but here the southern groups include the ≠Khomani, Nama, Karretjie (multiple language families) and the northern populations refer again to the !Xun and Ju/’hoansi. Thus, KhoeSan populations are not only strikingly isolated from other African populations but they appear geographically structured amongst themselves. To contrast this with Europeans, the ≠Khomani and the Ju/’hoansi may have diverged over 30,000 *ya* but live only 1,000 km apart, roughly the equivalent distance between Switzerland and Denmark whose populations have little genetic differentiation (Novembre et al. 2008). However, it is unclear how this ancient southern African divergence maps on to current linguistic and subsistence differences among populations, which may have emerged during the Holocene.

In particular, the genetic ancestry of the Khoe-speaking populations and specifically the Khoekhoe, (e.g. Nama) who practice sheep, goat and cattle pastoralism, remains a major open question. Archaeological data has been convened to argue for a demic migration of the Khoe from eastern African into southern Africa, but others have also argued that pastoralism represents cultural diffusion without significant population movement (Boonzaier 1996; K. C. MacDonald 2000; Robbins *et al.* 2005; Sadr 2008, 2015; Dunne *et al.* 2012; Pleurdeau *et al.* 2012; Jerardino *et al.* 2014). Lactase persistence alleles are present in KhoeSan groups, especially frequent in the Nama (20%), and clearly derive from eastern African pastoralist populations (Breton *et al.* 2014; Macholdt *et al.* 2014). This observation, in conjunction with other Y-chromosome and autosomal data (Henn *et al.* 2008; Pickrell *et al.* 2014), has been used to argue that pastoralism in southern Africa was another classic example of demic diffusion. However, the previous work is problematic in that it tended to focus on single loci [MCM6/LCT, Y-chromosome] subject to drift or selection. Estimates of eastern African autosomal ancestry in the KhoeSan remain minimal (<10%) and the distribution of ancestry informative markers is dispersed between both pastoralist and hunter-gatherer populations. Here, we present a comprehensive study of recent population structure in southern Africa and clarify fine-scale structure beyond “northern” and “southern” geographic descriptors. We then specifically test whether the Khoe-speaking Nama pastoralists derive their ancestry from eastern Africa, the northeastern Kalahari Basin, or far southern Africa. Our results suggest that ecological features of southern Africa, broadly speaking, are better explanatory features than either language, clinal geography or subsistence on its own.

## Results

To resolve fine-scale population structure and migration events in southern Africa, we generated genome-wide data from three South African populations. We genotyped ≠Khomani San (*n*=75), Nama (*n*=13) and South African Coloured (SAC) (*n*=25) individuals on the Illumina OmniExpress and OmniExpressPlus SNP array platforms. Sampling locations are listed in Table S1, in addition to language groupings and subsistence strategies. These data were merged with HapMap3 (joint Illumina Human1M and Affymetrix SNP 6.0) (Consortium 2010), HGDP (Illumina 650Y) data (Li *et al.* 2008) and Petersen et al. Illumina HumanOmni1-Quad (Petersen *et al.* 2013), resulting in an intersection of ˜320k SNPs for 852 individuals from 21 populations. In addition, we used the Affymetrix Human Origins SNP Array generated as part of Pickrell et al. (Pickrell *et al.* 2012) and Lazaridis et al. (Lazaridis *et al.* 2014), including *n*=9 ≠Khomani San individuals from our collection and encompassing over 396 individuals from 33 populations. Whole mitochondrial genomes were generated from off-target reads from exome- and Y chromosome-capture short read Illumina sequencing. Reads were mapped to GRCh37, which uses the revised Cambridge reference sequence (rCRS). Only individuals with greater than 7x haploid coverage were included in the analysis: ≠Khomani San (*n*=64) and Nama (*n*=31); haplogroup frequencies were corrected for pedigree structure (Table S2). In this study, we address population structure among southern African KhoeSan, the genetic affinity of the Khoe, and how pastoralism diffused into southern Africa.

### Population Structure in Southern African KhoeSan Populations

We first tested whether southern African populations conform to an isolation-by-distance model, or whether there is strong heterogeneity among populations relative to geographic distance. Using twenty-two southern African populations (with 560k SNPs from Affymetrix HumanOrigins array), we implemented the spatially explicit program EEMS (Petkova *et al.* 2016) to test for effective migration patterns across the region. We observe a higher effective migration rate (*m*) in the central Kalahari basin relative to a lower migration rate that forms a rim around the Kalahari Desert (Figure 1). A second resistance band stretches across northern Namibia, indicating higher gene flow above northern Namibia, Angola and southern Zambia. Differences in effective migration rates can result from differences in effective population sizes. For example, a larger effective population size can result in higher effective migration rates, relative to neighboring demes with smaller *N_e_*’s. The higher *m* in the central Kalahari Basin, relative to the rim, could result from either a larger *N_e_* relative to Kalahari rim populations or simply higher migration among groups in a similar ecological area.

**Figure 1:**
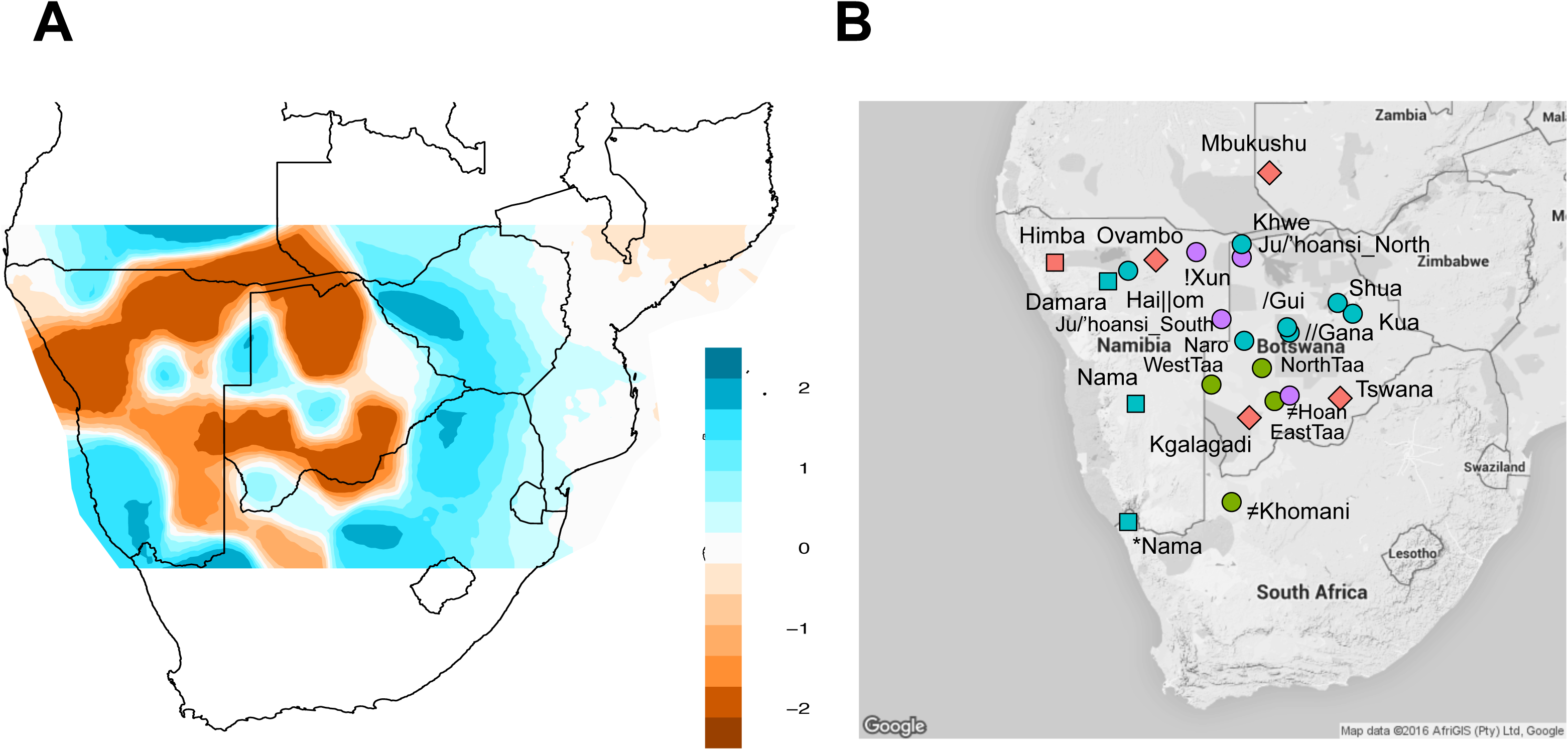
Effective migration rates among 22 southern African populations. A) Using southern African samples from the Affymetrix HumanOrigins dataset, we estimated effective migration rates among populations using EEMS. White indicates the mean expected migration rate across the dataset, while blue indicates X-fold increase in migration among demes, and brown indicates decreased migration among demes (e.g. population structure). Effective migration rates, *e_m_*, are plotted on a log-scale as in Petkova et al (2016). Hence, −1 *e_m_* would indicate 10-fold decrease in the migration rate relative to the expected rate among all demes accounting for geographic distance. These results demonstrate that southern Africa is a heterogeneous environment with barriers to gene flow in northwest Namibia and the Kalahari rim, but increased gene flow within the Kalahari Basin. The grid of plotted demes was restricted to prevent unwanted extrapolation to poorly sampled areas. B) The topographic map indicates the subsistence strategy and language of each population sample. Colors represent language families: green= Tuu speakers, red= Niger-Congo speakers, blue= Khoe speakers and purple= Kx’a speakers. Shapes represent subsistence strategies: circle= hunter-gatherers, square= pastoralists and diamond= agropastoralists. *Nama indicates a new, second Nama sample from South Africa, which was only included in Illumina SNP array analyses.

We then tested whether heterogeneity in population structure could be mapped to distinct genetic ancestries. Unsupervised population structure analysis identifies 5 distinct, spatially organized ancestries among the sampled twenty-two southern African populations. These ancestries were inferred from the Affymetrix Human Origins dataset using ADMIXTURE (Figure S1) (Alexander *et al.* 2009). Multi-modality per *k* value was assessed using *pong* (Behr *et al.* 2015) and results from k=10 are discussed below (6/10 runs assigned to the major mode, 3/10 other runs involved cluster switching only within East Africa). Visualization of these ancestries according to geographic sampling location specifically demonstrates fine-scale structure in and around the Kalahari Desert (Figure 2). While prior studies have argued for a northern versus southern divergence of KhoeSan populations (Pickrell *et al.* 2012; Schlebusch *et al.* 2012; Schlebusch and Soodyall 2012; Barbieri *et al.* 2013, 2014), the structure inferred from our dataset indicates a more geographically complex pattern of divergence and gene flow. Even recent migration events into southern Africa remain structured, consistent with ecological boundaries to gene flow (see below). The distribution of the five ancestries corresponds to: a northern Kalahari ancestry, central Kalahari ancestry, circum-Kalahari ancestry, a northwestern Namibian savannah ancestry and ancestry from eastern Bantu-speakers (Figure 2). This geographic patterning does not neatly correspond to linguistic or subsistence categories, in contrast to previous discussions (Pickrell *et al.* 2012; Schlebusch *et al.* 2012; Barbieri *et al.* 2014).

**Figure 2:**
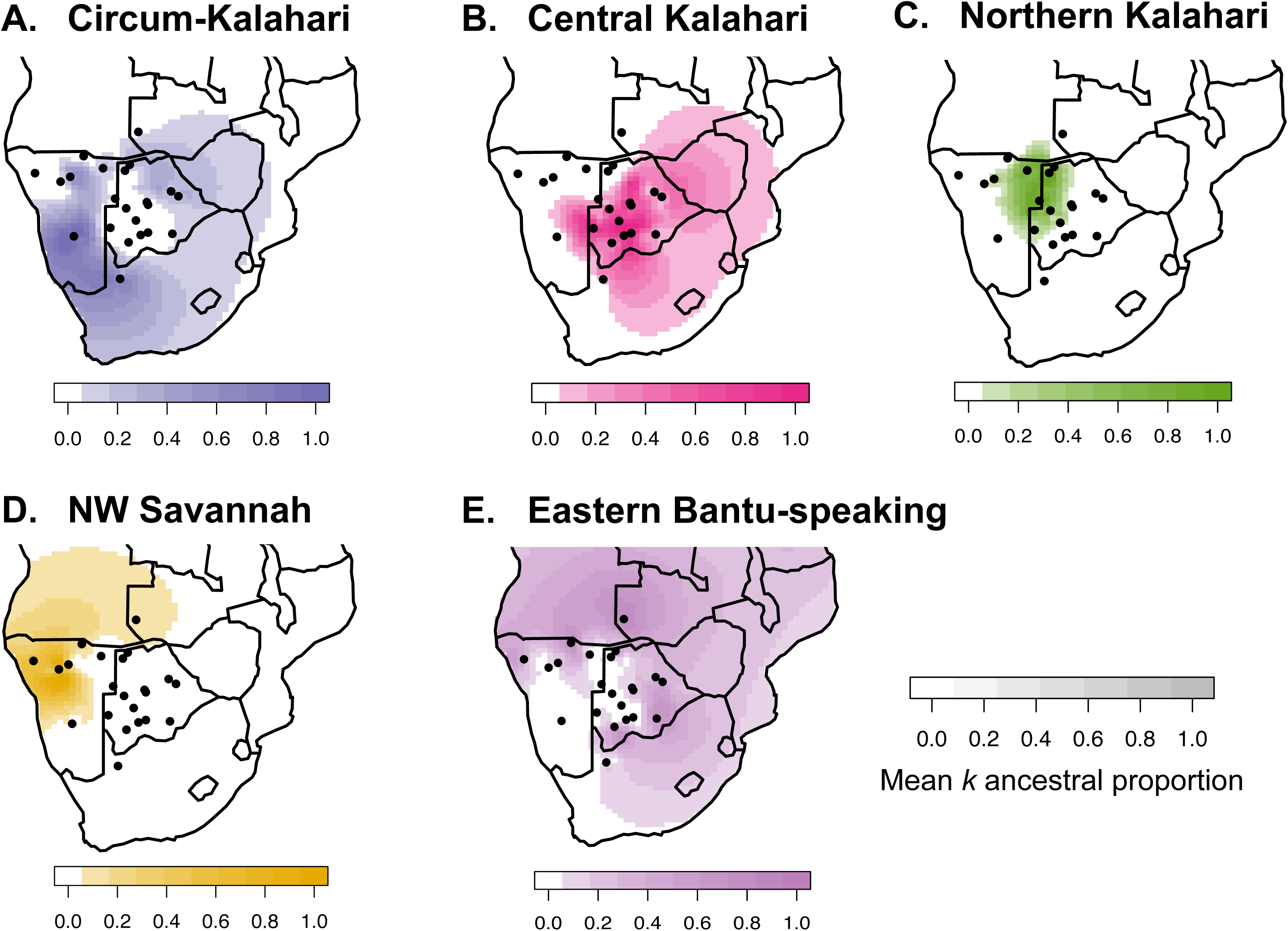
Five spatially distinct ancestries indicate deep population structure in southern Africa. Using global ancestry proportions inferred from ADMIXTURE *k*=10, we plot the mean ancestry for each population in southern Africa. The 5 most common ancestries in southern Africa, from the Affymetrix HumanOrigins dataset, are shown separately in panels A)-E). The X and Y axes for each map correspond to latitude and longitude, respectively. Black dots represent the sampling location of populations in southern Africa. The 3^rd^ dimension in each map [depth of color] represents the mean ancestry proportion for each group for a given *k* ancestry, calculated from ADMIXTURE using unrelated individuals, and indicated in the color keys as 0% to 100% for five specific *k* ancestries. Surface plots of the ancestry proportions were interpolated across the African continent.

The northern Kalahari ancestry is the most defined of these ancestries, encompassing several forager populations such as the Ju/’hoansi, !Xun, Khwe, Naro and to a lesser extent the Khoekhoe-speaking Hai||om. While these populations are among the best-studied KhoeSan in anthropological texts with particular reference to cultural similarities (Dornan 1925; Bleek 1928; Schapera 1934; Barnard 1992a), they represent only a fraction of the diversity among Khoisan-speaking populations. We note that this cluster includes Kx’a (Juu), Khoe-Kwadi and Khoekhoe speakers, suggesting that language interacts in a complex fashion with other factors such as subsistence strategy and ecology. The Hai∥om are thought to have shifted to speaking Khoekhoe from an ancestral Juu-based language (Barnard 1992a). The second, central Kalahari ancestry occupies a larger geographical area throughout the Kalahari basin, with its highest frequency among the Taa-speakers: G|ui, G∥ana, ≠Hoan and Naro. This ancestry spans all three Khoisan language families (Table S1), at considerable frequency in each; all are primarily foragers.

The third ancestry cluster is represented by southern KhoeSan populations distributed along the rim of the Kalahari Desert (Figure 2) – referred to here as the “circum-Kalahari ancestry”. The circum-Kalahari ancestry is at its highest frequency in the Nama and ≠Khomani (see also Figure S2), with significant representation in the Hai∥om, Khwe, !Xun and Shua. This ancestry spans all linguistic and subsistence strategies. We propose that the circum-Kalahari is better explained by ecology than alternative factors such as language or recent migration. Specifically, we find the Kalahari Desert is an ecological boundary to gene flow (Figures 1,2). The circum-Kalahari ancestry is not easily explained by a pastoralist Khoekhoe dispersal. This spatially distinct ancestry is common in both forager and pastoralist groups, indeed all of the circum-Kalahari populations were historically foragers (except for the Nama). Therefore, to support a Khoekhoe dispersal model, we would have to posit an adoption of pastoralism by a northeastern group, leading to demic expansion around the Kalahari, with subsequent reversion to foraging in the majority of the circum-Kalahari groups; this scenario seems unlikely (but see Smith (2014) for additional discussion).

Finally, our analysis reveals two additional ancestries outside of the greater Kalahari Basin: one ancestry composed of Bantu-speakers, frequent to the north, east and southeast of the Kalahari; and a second composed of Himba, Ovambo, and Damara ancestry in northwestern Namibia distributed throughout the mopane savannah. Interestingly, the Damara are a Khoekhoe-speaking population of former foragers (later in servitude to the Nama pastoralists) whose ancestry has been unclear (*see below*).

We used our data and the Affymetrix HumanOrigins dataset containing the greatest number of KhoeSan populations to date, to test whether language or geography better explains genetic distance (see language families and subsistence strategies in Table S1). The genetic data were compared to a phonemic distance matrix (Jaccard 1908) as well as geographic distances between each population (Table S3). In order to test whether genetic distance (F_st_) was associated with geography or language, we performed a partial Mantel test for the relationship between F_st_ and language (Creanza *et al.* 2015) accounting for geographic distance among 11 KhoeSan populations. This result was not significant (r=0.06, p=0.30). Although an association between F_st_ and geographical distance within Africa has been documented (Ramachandran *et al.* 2005; Tishkoff *et al.* 2009; Creanza *et al.* 2015), a Mantel test for the relationship between F_st_ and pairwise geographic distance in our dataset was also null (r=0.021, p=0.38) reflecting the non-linear aspect of shared ancestry in southern Africa as seen in Figures 1 and 2.

Spatially distinct ancestries are also supported by principal components analysis (PCA) (Figure 3 and S3). The KhoeSan anchor one end of PC1 opposite to Eurasians. PC2 separates other African populations from the KhoeSan, including western Africans, as well as central and eastern African hunter-gatherers. PC3 separates the Ju/’hoansi and !Xun (northern Kalahari) from ≠Hoan, Taa-speakers and Khoe-speakers, with other KhoeSan populations intermediate. PC3 and PC4 suggest that the present language distribution may reflect recent language transitions, as genetic ancestry and linguistic structure do not neatly map onto each other (**Figure S4**). For example, the ≠Hoan currently speak at Kx’a language but are genetically distinct from other northern Kalahari Kx’a speakers; rather, they appear to be more genetically similar to southern Kalahari Taa-speakers who cluster together. We suggest that the patterns observed here are better explained by ecogeographic patterns than either language or subsistence alone (**Figure S5**). Specifically, PC3 discriminates northern versus southern Kalahari ancestry (see below). PC4 discriminates western and eastern non-KhoeSan ancestry derived from Bantu-speakers or other populations. Finally, the intermediate position of the Nama, ≠Khomani and Hai‖om on PC3 and PC4 is neither linguistic- nor subsistence-based, but represents a non-linear circum-Kalahari component featured in Figure 2.

**Figure 3:**
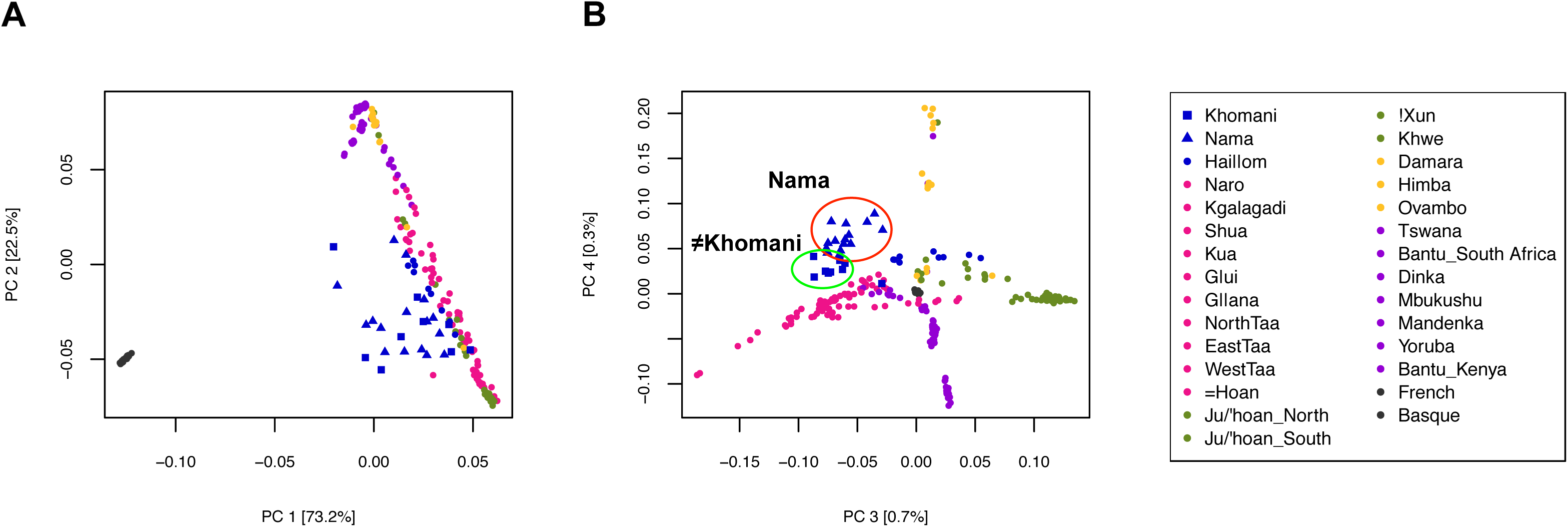
Clustering of KhoeSan populations and fine-scale population structure between the Nama and **≠**Khomani San. A PCA of the Affymetrix Human Origins dataset depicts the clustering of unrelated individuals based on the variation seen in the dataset. Colors mimic similar major ancestry colors as shown in Figure 2. Yellow denotes populations with majority northwestern Namibian ancestry; purple denotes populations with majority Bantu-speaking ancestry; pink indicates southern Kalahari majority ancestry, green indicates northern Kalahari majority ancestry and blue indicates circum-Kalahari ancestry. The red and green circles denote the fine-scale separation of the Nama and **≠**Khomani populations [specified by triangles and squares, respectively]. Note that these colored ancestries and the PCs do not map onto subsistence neatly (Figure S5).

### A Divergent Southern KhoeSan Ancestry

This separation of northern (Ju/’hoansi) and southern (Taa and Khoe speakers) KhoeSan populations has been observed by Schlebusch et al. (Schlebusch *et al.* 2012) and Pickrell et al. (Pickrell *et al.* 2012). We estimate that this trans-Kalahari genetic differentiation from the inferred ancestral allele frequencies (Figure S2) is substantial (F_ST_ = 0.05). We verify this divergence between the northern Kx’a-speakers and the shared Nama and ≠Khomani ancestry in a new, second sample of Nama, from South Africa rather than central Namibia (Table S1, Figure S3). This southern KhoeSan ancestry is also present in admixed Bantu-speaking populations from South Africa (e.g. amaXhosa) as well as the admixed Western Cape SAC populations (de Wit *et al.* 2010), supporting a hypothesis of distinct *southern*-specific KhoeSan ancestry (Figure S1, S2) shared between indigenous and admixed groups.

Mitochondrial data support this concept of a *southern*-specific KhoeSan ancestry (Schlebusch *et al.* 2013; Barbieri *et al.* 2013). Both mtDNA haplogroups L0d and L0k are at high frequency in northern KhoeSan populations (Behar *et al.* 2008), but L0k is absent in our sample of the Nama [*n*=31] and there is only one ≠Khomani individual [n=64] with L0k (1.56%) (Table 1). L0d dominates the haplogroup distribution for both the Nama and ≠Khomani (84% and 91% respectively), with L0d2a especially common in both. L0d2a, inferred to have originated in southern Africa, was also previously found at high frequencies in the Karretjie people further south in the central Karoo of South Africa, as well as the SAC population in the Western Cape (Quintana-Murci *et al.* 2010; Schlebusch *et al.* 2013). L0d2b is also common in the Nama (16%).

**Table 1:** Mitochondrial DNA haplogroup frequencies of the Nama and **≠**Khomani

|  |  | #Khomani San |  |  | Nama |  |  |
| --- | --- | --- | --- | --- | --- | --- | --- |
| Haplogroup |  | n | Frequency |  | n | Frequency |  |
| L0d | L0d1a | 8 | 12.50% | 90.63% | 3 | 9.68% | 83.87% |
|  | L0d1a1 | 2 | 3.13% |  | 3 | 9.68% |  |
|  | L0d1b | 4 | 6.25% |  | 3 | 9.68% |  |
|  | L0d1b1 | 9 | 14.06% |  | 2 | 6.45% |  |
|  | L0d1b2 | 2 | 3.13% |  | 0 |  |  |
|  | L0d1c1 | 1 | 1.56% |  | 1 | 3.23% |  |
|  | L0d1c1a | 1 | 1.56% |  | 1 | 3.23% |  |
|  | L0d2a | 25 | 39.06% |  | 3 | 9.68% |  |
|  | L0d2a1 | 0 |  |  | 2 | 6.45% |  |
|  | L0d2b | 0 |  |  | 5 | 16.13% |  |
|  | L0d2c | 5 | 7.81% |  | 2 | 6.45% |  |
|  | L0d3 | 1 | 1.56% |  | 1 | 3.23% |  |
| L0f1 |  | 1 | 1.56% |  | 0 |  |  |
| L0k1 |  | 1 | 1.56% |  | 0 |  |  |
| L3'4 |  | 0 |  |  | 1 | 3.23% |  |
| L3d3a1a |  | 0 |  |  | 1 | 3.23% |  |
| L3e1a2 |  | 1 | 1.56% |  | 0 |  |  |
| L4b2a2 |  | 1 | 1.56% |  | 1 | 3.23% |  |
| L5c |  | 0 |  |  | 1 | 3.23% |  |
| M36 | M36 | 1 | 1.56% | 3.13% | 0 |  |  |
|  | M36d1 | 1 | 1.56% |  | 0 |  |  |
| M7c3c |  | 0 |  |  | 1 | 3.23% |  |
| Total (n) |  | 64 | 100% |  | 31 | 100% |  |

### Minimal Population Structure Between the Nama and ≠Khomani

The ≠Khomani San are a N|u-speaking (!Ui classified language) former hunter-gatherer population that inhabit the southern Kalahari Desert in South Africa, bordering on Botswana and Namibia. The Nama, currently a primarily caprid pastoralist population, live in the Richtersveld along the northwestern coast of South Africa and up into Namibia. The ancestral geographic origin of the Nama has been widely contested over a number of years (Nurse and Jenkins 1977; Barnard 1992b; Boonzaier 1996), but a leading hypothesis suggests that they originated further north in Botswana/Zambia and migrated into South Africa and Namibia approximately 2,000 years ago (Nurse and Jenkins 1977; Barnard 1992b; Boonzaier 1996; Pickrell *et al.* 2012). The Nama and N|u languages are in distinct, separate Khoisan language families (Khoe and Tuu [!Ui-Taa], respectively) and these groups historically utilized different subsistence strategies. For this reason, we hypothesized that there would be strong population structure between the two populations.

Our global ancestry results, inferred from ADMIXTURE, show minimal population structure between the Nama and ≠Khomani San in terms of their southern KhoeSan ancestry. The ≠Khomani share ˜10% of their ancestry with the Botswana KhoeSan populations (Figure S1, S3), consistent with their closer proximity to the southern Botswana populations (Taa-speakers !Xo and ≠Hoan). Principal components analysis reveals a degree of fine-scale population structure between the Nama and ≠Khomani, with each population forming its own distinct cluster at PC4, partly due to the increase in Damara ancestry in the Nama (Figure 3b, Figure S1), but the two groups are clearly proximal. This increase in Damara ancestry (as depicted from *k*=9 in all modes of Figure S1) is likely due to integration of the Damara people as clients of the Nama over multiple generations. However, our second sample of Nama from South Africa do not harbor significant western African ancestry, suggesting heterogeneity in the Damara component (Figure S2).

### Recent Patterns of Admixture in South Africa

Two Bantu-speaking, spatially distinct ancestries are present in southern Africa. The first is rooted in the Ovambo and Himba in northwestern Namibia; the other reflects gene flow from Bantu-speaking ancestry present in the east (Figure 2). We estimated the time intervals for admixture events into the southern KhoeSan via analysis of the distribution of local ancestry segments using RFMix (Maples *et al.* 2013) and TRACTs (Gravel 2012) for the **≠**Khomani OmniExpress dataset (n=59 unrelated individuals) (Figure 4, Table S2). The highest likelihood model suggests that there were 3 gene flow events. Approximately 14 generations ago (˜443-473 years ago assuming a generation time of 30 years and accounting for the age of our sampled individuals), the **≠**Khomani population received gene flow from a Bantu-speaking group, represented here by the Kenyan Luhya. Our results are consistent with Pickrell et al. (2012) who found that the southern Kalahari Taa-speakers were the last to interact with the expanding Bantu-speakers about 10-15 generations ago. Subsequently, this event was followed by admixture with Europeans between 6 and 7 generations ago (˜233-263 years ago), after the arrival of the Dutch in the Cape and the resulting migrations of “trekboers” (nomadic pastoralists of Dutch, French and German descent) from the Cape into the South African interior. Lastly, we find a recent pulse of primarily KhoeSan ancestry 4-5 generations ago (˜173-203 years ago). This event could be explained by gene flow into the ≠Khomani from another KhoeSan group, potentially as groups shifted local ranges in response to the expansion of European farmers in the Northern Cape, or other population movements in southern Namibia or Botswana.

**Figure 4:**
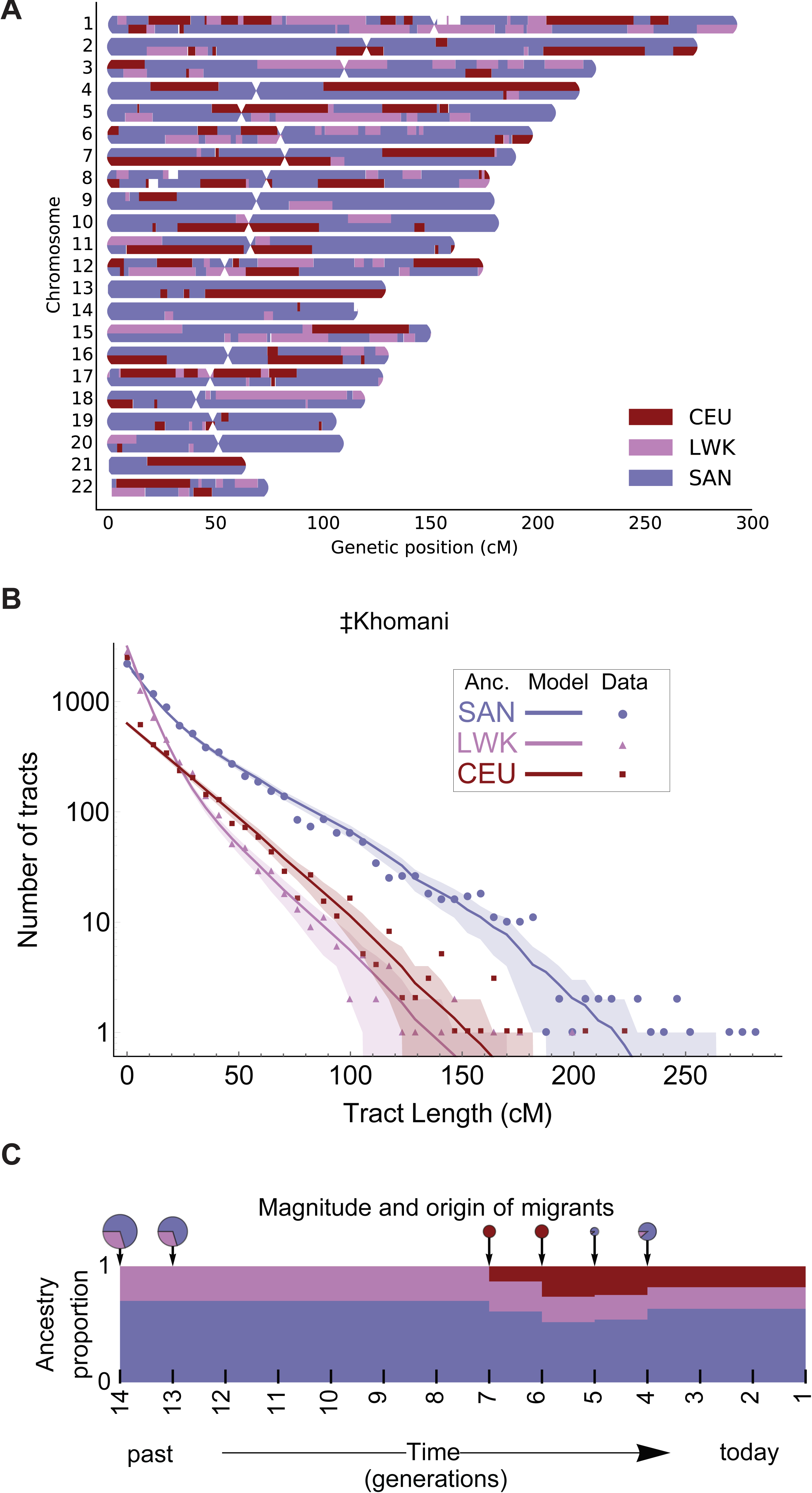
Demographic reconstruction of recent admixture in the **≠**Khomani San using local ancestry. A) A local ancestry karyogram for a representative 3-way admixed ≠Khomani San individual was constructed using RFMix. Haplotypes for admixed individuals were assigned to one of three possible ancestries: SAN [Namibian San], LWK [Bantu-speaking Luhya from Kenya], CEU [Central Europeans]. UNK indicates “unknown” ancestry (*Methods*). B) We employed Markov models implemented in *TRACTs* to test multiple demographic models and assess the best fit to the observed ≠Khomani haplotype distributions. Local ancestry tract lengths were inferred as in panel A). The tract length distribution for each ancestry across all individuals was used to estimate migration time (generations ago), volume of migrants, and ancestry proportions over time. Colored dots show the observed distribution of ancestry tracts for each ancestry, solid lines show the best fit from the most likely model, and shaded areas indicate confidence intervals corresponding to ± 1 standard deviation.

We also considered the impact of recent immigration into indigenous South Africans, derived from non-African source populations. The South African Coloured (SAC) populations are a five way admixed population, deriving ancestries from Europe, eastern African, KhoeSan, and Asian populations (de Wit *et al.* 2010). This unique, admixed ethnic population was founded by the Dutch who settled on the southern tip of South Africa by the 17^th^ century and the importation of slaves from Indonesia, Bengal, India and Madagascar. However, within the SAC, strong differences in ancestry and admixture proportions are observed between different districts within Cape Town, the Eastern Cape and the Northern Cape Provinces. South African Coloured individuals from the Northern Cape, where historically there was a greater concentration of European settlement (Theal 1887), have higher European ancestry. The SAC individuals from the Eastern Cape, which is the homeland of the Bantu-speaking Xhosa populations, have relatively more ancestry from Bantu-speaking populations (Figure S2). The “ColouredD6” population is from an area in Cape Town called District 6. Historically, this was a district where the slaves and political exiles from present day Indonesia resided, and many who were from Madagascar and India based on written documentation (du Plessis 1947). The SAC D6 population consequently has a noticeable increase in south/eastern Asian ancestry represented by the Pathan and Han Chinese populations in our dataset (Figure S2).

This south/eastern Asian ancestry is not confined to the SAC population, as attested by the presence of the M36 mitochondrial haplogroup. The M36 haplogroup (South Indian/Dravidian in origin) is present in two out of 64 ≠Khomani San matrilineages, (**Table 1**). The presence of M36 is likely derived from slaves of South Asian origin who escaped from Cape Town or the surrounding farms and dispersed into the northwestern region of South Africa. In addition, we observe one M7c3c lineage in the Nama (**Table 1**), which traces back to southeastern Asia but has been implicated in the Austronesian expansion of Polynesian speakers into Oceania (Kayser 2010; Delfin *et al.* 2012) and Madagascar (Poetsch *et al.* 2013). The importation of Malagasy slaves to Cape Town may best explain the observation of M7c3c in the Nama.

## Discussion

The KhoeSan are distinguished by their unique phenotype(s), genetic divergence, click languages and hunter-gatherer subsistence strategy compared to other African populations; classifications of the many KhoeSan ethnic groups have primarily relied on language or subsistence strategy. Here, we generate additional genome-wide data from 3 South African populations and explore patterns of fine-scale population structure among 22 southern African groups. We find that complex geographic or “ecological” information is likely a better explanatory variable for genetic ancestry than language or subsistence. We identify 5 primary ancestries in southern Africans, each localized to a specific geographic region (Figure 2). In particular, we examined the “circum-Kalahari” which appears as a ring around the Kalahari Desert and accounts for the primary ancestry of the Nama, representative of the Khoekhoe-speaking pastoralists.

We observe striking ecogeographic population structure associated with the Kalahari Desert. There are two distinct ancestries segregating within the Kalahari Desert KhoeSan populations, described here as northern Kalahari and central Kalahari ancestries. Analyses of migration rates across the 22 populations indicate particularly high migration within the Kalahari Desert. This may indicate a larger effective population size for the two desert ancestries or extensive migration related to shifting ranges in response to climatic and ecological changes over time. It is worth noting that the northern Kalahari formerly supported an extensive lake (i.e. Makgadikgadi) just before and after the Last Glacial Maximum, as well as the presence of the Okavango Delta and associated river systems; archeological data may suggest high population density nearby the pans, although this likely predates the genetic structure we observe today (Burrough 2016; Robbins *et al.* 2016). Our lack of samples outside of Botswana, Namibia and northern South Africa prevent precise inference of *m* in Zambia, Limpopo, and Mozambique; but Figure 2 indicates recent extensive gene flow in the east, consistent with the expansion of Bantu-speaking agriculturalists into eastern grasslands and coastal forests. Additionally, we find a separate ancestry segregating in the far western border of Namibia and Angola, particularly frequent in the Damara and Himba, and to a lesser extent in the Ovambo and Mbukushu. This intersection of steppe and savannah along the Kunene may have facilitated recent settlement of the area during the past 500 years by Bantu-speaking pastoralists, but it is noteworthy that little Kalahari KhoeSan ancestry persists in these populations. Rather, the Damara (currently Nama-speaking) or related hunter-gatherers may have been formerly more widespread in this area and subsequently absorbed into the western Bantu-speaking pastoralists.

The practice of sheep, goat and cattle pastoralism in Africa is widespread. Within KhoeSan populations, pastoralist communities are limited to the Khoekhoe-speaking populations. Earlier hypotheses proposed that the Khoe-speaking pastoralists derived from a population originating outside of southern Africa. However, more recent genetic work supports a model of autochthonous Khoe ancestry influenced by either demic or cultural diffusion of pastoralism from East Africa ˜2,500 years ago (Pleurdeau *et al.* 2012; Pickrell *et al.* 2014). For example, the presence of lactase persistence alleles in southern Africa indicates contact between East African herders and populations in south-central Africa, with subsequent migration into Namibia (Breton *et al.* 2014). This scenario is also supported by Y-chromosomal analysis that indicates a direct interaction between eastern African populations and southern African populations approximately 2,000 years ago (Henn *et al.* 2008). However, in both cases (i.e. MCM6/LCT and Y-chr M293), the frequency of the eastern African alleles is low in southern Africa and occurs in both pastoralist and hunter-gatherer populations. A simple model of eastern African demic diffusion into south-central Africa, leading to the adoption of pastoralism and a Khoekhoe population expansion from this area cannot be inferred from the genetic data.

Our samples from the Khoekhoe-speaking Nama pastoralists demonstrate that their *primary* ancestry is shared with other far southern non-pastoralist KhoeSan such as the ≠Khomani San and the Karretjie (see also Schlebusch *et al.* 2011). mtDNA also suggests that the Nama display a haplogroup frequency distribution more similar to KhoeSan south of the Kalahari than to any other population in south-central Africa. Our results indicate that the majority of the Nama ancestry has likely been present in far southern Africa for longer than previously assumed, *rather* than resulting from a recent migration from further north in Botswana where other Khoe-speakers live. The only other Khoekhoe-speaking population in our dataset is the Hai‖om who share approximately 50% of the circum-Kalahari ancestry with the Nama and ≠Khomani, but are foragers rather than pastoralists. We conclude that Khoekhoe-speaking populations share a circum-Kalahari genetic ancestry with a variety of other Khoe-speaking forager populations in addition to the !Xun, Karretjie and ≠Khomani (Figures 1 and 2). This ancestry is divergent from central and northern Kalahari ancestries, arguing *against* a major demic expansion of Khoekhoe pastoralists from northern Botswana into South Africa. Rather, in this region, cultural transfer likely played a more important role in the diffusion of pastoralism. Of course, a demic expansion of the Khoekhoe *within* a more limited region of Namibia and South Africa may still be have occurred – but geneticists currently lack representative DNA samples from many of the now “Coloured” interior populations which may carry Khoekhoe ancestry.

This is an unusual case of cultural transmission (Jerardino *et al.* 2014). Other prehistoric economic transitions have been shown to be largely driven by demic diffusion (Gignoux *et al.* 2011; Fort 2012; Skoglund *et al.* 2014; Lazaridis *et al.* 2014; Malmström *et al.* 2015). Recent analysis of Europe provides a case study of demic diffusion, which appears far more complex than initially hypothesized. The initial spread of Near Eastern agriculturalists into southern Europe clearly replaced or integrated many of the autochthonous hunter-gatherer communities. Even isolated populations such as the Basque have been shown to derive much of their ancestry from Near Eastern agriculturalists (Skoglund *et al.* 2014). The early demic diffusion of agriculture exhibits a strong south to north cline across Europe, reflecting the integration of hunter-gatherers into composite southern agriculturalist populations which then expanded northward with mixed ancestry (Sikora *et al.* 2014). The cline of the early Near Eastern Neolithic ancestry becomes progressively diluted in far northern European populations. In contrast, we see little evidence of a clear eastern African ancestry cline within southern African KhoeSan; nor is the putative “Khoe” ancestry identified in the Nama of eastern African origin or even of clear origin from northeastern Botswana where initial pastoralist contact presumably occurred.

However, the transfer of pastoralism from eastern to southern Africa itself was not purely cultural (see above). We also report here the presence of mitochondrial L4b2 that supports limited gene flow from eastern Africa, approximately during the same timeframe as the pastoralist diffusion. L4b2, formerly known as L3g or L4g, is a mtDNA haplogroup historically found at a high frequency in eastern Africa, in addition to the Arabian Peninsula. L4b2 is at high frequency specifically in click-speaking populations such as the Hadza and Sandawe in Tanzania (sometimes described as ‘Khoisan-speaking’) (Knight *et al.* 2003). Nearly 60% of the Hadza population and 48% of Sandawe belong to L4b2 (Tishkoff *et al.* 2007). Even though both Tanzanian click-speaking groups and the southern African KhoeSan share some linguistic similarities and a hunter-gatherer lifestyle, they have been isolated from each other over the past 35*ky* (Tishkoff *et al.* 2007). The L4b2a2 haplogroup is present at a low frequency in both the Nama and ≠Khomani San, observed in one matriline in each population (**Table 1**). L4b2 was also formerly reported in the SAC population (0.89%) (Quintana-Murci *et al.* 2010) but has not been discussed in the literature. We identified several additional southern L4b2 haplotypes from whole mtDNA genomes deposited in public databases (Behar *et al.* 2008; Barbieri *et al.* 2013) and analyzed these samples together with all L4b2 individuals available in NCBI. Median-joining phylogenetic network analysis of the mtDNA haplogroup, L4b2, supports the hypothesis that there was gene flow from eastern Africans to southern African KhoeSan groups. As shown in Figure 5 (and in more detail in Figure S6), southern African individuals branch off in a single lineage from eastern African populations in this network (Salas *et al.* 2002; Tishkoff *et al.* 2007; Gonder *et al.* 2007). The mitochondrial network suggests a recent migratory scenario (estimated to be < 5,000 years before present), though the source of this gene flow, whether from eastern African click-speaking groups or others, remains unclear (Pickrell *et al.* 2014).

**Figure 5:**
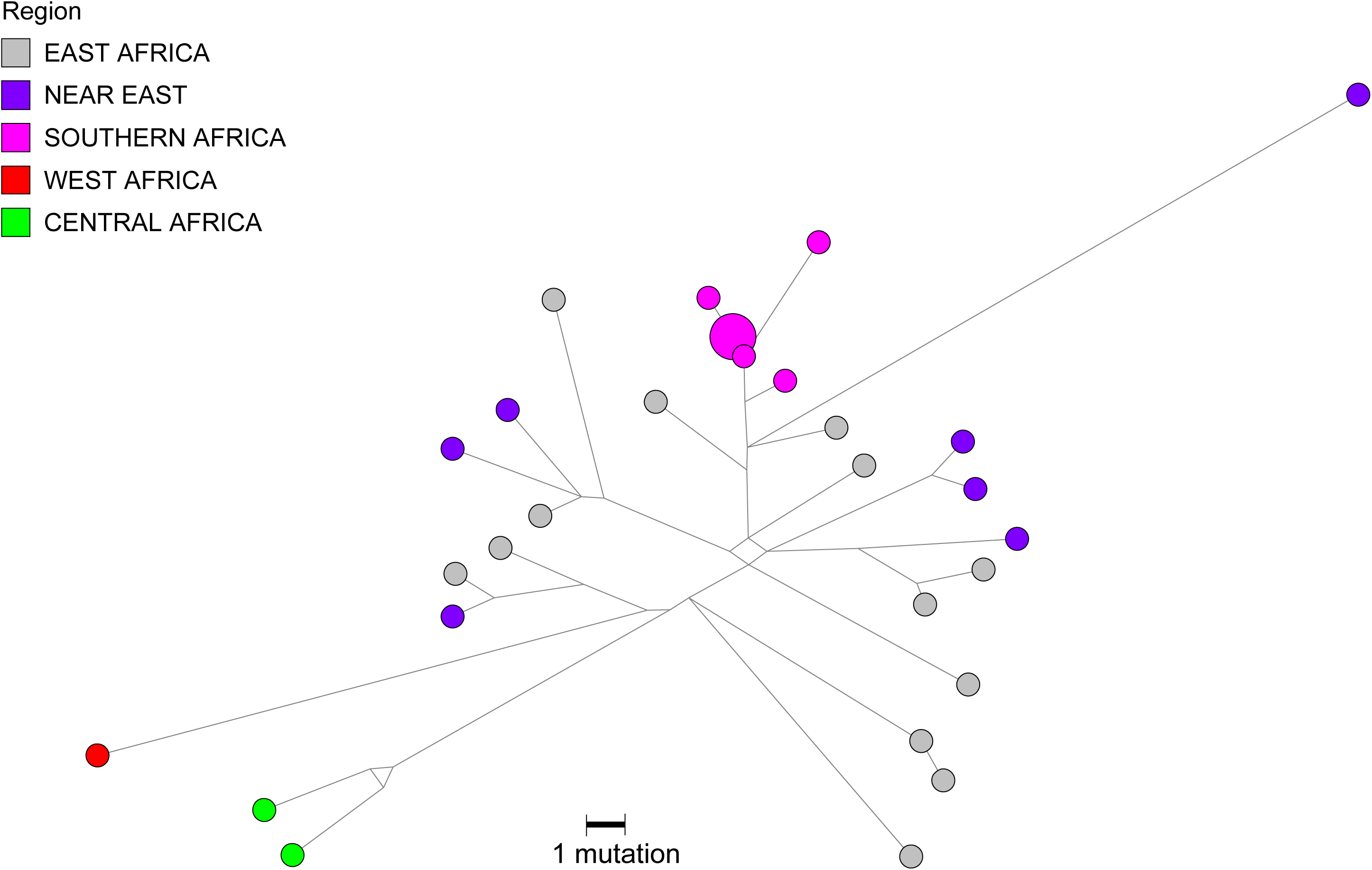
L4b2 mtDNA haplogroup network. New L4b2 mitochondrial genomes from **≠** Khomani and Nama individuals, indicated in pink as “Southern Africa”, were analyzed together with publically available L4b2 mtDNA genomes from NCBI (as outlined in the *Supplementary Methods*). All individuals were assigned to mtDNA haplogroups using *haplogrep* and the haplotypes were plotted using Network Publisher.

## Conclusion

Analysis of 22 southern African populations reveals that fine-scale population structure corresponds better with ecological rather than linguistic or subsistence categories. The Nama pastoralists are autochthonous to far southwestern Africa, rather than representing a recent population movement from further north. We find that the KhoeSan ancestry remains highly structured across southern Africa and suggests that cultural diffusion likely played the key role in adoption of pastoralism.

## Materials and Methods

### Sample collection and ethical approval

DNA samples from the Nama, ≠Khomani San and South African Coloured populations were collected with written informed consent and approval of the Human Research Ethics Committee of Stellenbosch University (N11/07/210), South Africa, and Stanford University (Protocol 13829), USA. Community level results were returned to the communities in 2015 prior to publication. A contract for this project was approved by WIMSA (ongoing).

### Autosomal data and genotyping platforms

A) ˜565,000 SNP Affymetrix Human Origins SNP array dataset from Pickrell et al. (Pickrell *et al.* 2012), Lazaridis et al. (Lazaridis *et al.* 2014) and additional ≠Khomani San and Hadza individuals from our collections: 33 populations and 396 individuals. B) ˜320,000 SNP array dataset from the intersection of HGDP (Illumina 650Y) (Li *et al.* 2008), HapMap3 (joint Illumina Human 1M and Affymetrix SNP 6.0), Illumina OmniExpressPlus and OmniExpress SNP array platforms as well as the dataset from Petersen et al. (Petersen *et al.* 2013): 21 populations and 852 individuals.

### Population structure

ADMIXTURE (Alexander *et al.* 2009) was used to estimate the ancestry proportions in a model-based approach. Iterations through various *k* values are necessary. The *k* value is an estimate of the number of original ancestral populations. Cross-validation (CV) was performed by ADMIXTURE and these values were plotted to acquire the *k* value that was the most stable. Depiction of the Q matrix was performed in R. Ten iterations were performed for each *k* value with ten random seeds. Iterations were grouped according to admixture patterns to identify the major and minor modes by pong (Behr *et al. 2015*). These Q matrixes from ADMIXTURE, as well as longitude and latitude coordinates for each population were adjusted to the required format for use in an R script supplied by Prof. Ryan Raaum to generate the surface maps (Figure 2).

### EEMS analysis

EEMS (Petkova *et al.* 2016) was run on the Affymetrix Human Origins dataset. Genetic dissimilarities were calculated using the bed2diffs script and EEMS was run using the runeems_snps version of the program. A grid is constructed so as to house all demes in the data provided. Each individual is assigned to a specific deme. Using a stepping stone model, migration rates between demes are calculated. Genetic dissimilarities are calculated fitting an “isolation by distance model”. In order for the MCMC iterations to converge, the number of MCMC iterations, burn iterations and thin iterations were increased. The other parameters were optimized as per the manual’s recommendations, i.e. diversity and migration parameters were adjusted so as to produce 20-30% acceptance rates. The PopGPlot R package was used to visualize the data.

### Association between F_st_, geography and language

A Mantel test (Fst and geographic distance) and a partial Mantel test (F_st_ and language, accounting for geographic distance) was performed using the vegan package in R. Geographic distances (in kilometers) between populations were calculated using latitude and longitude values as tabulated in Table S1. Weir and Cockerham genetic distances (F_st_) were calculated from allele frequencies estimated with vcftools (Danecek *et al.* 2011). A Jaccard phonemic distance matrix was used as formulated in Creanza et al. (Creanza *et al.* 2015). Populations included in the analysis were the Nama, ≠Khomani, East Taa, West Taa, Naro, G|ui, G∥ana, Shua, Kua, !Xuun and Khwe.

### mtDNA Network

We utilized Network (ver. 4.6, copy righted by Fluxus Technology Ltd.), for a median-joining phylogenetic network analysis in order to produce Figures 5 and S6. Network Publisher (ver. 2.0.0.1, copy righted by Fluxus Technology Ltd.) was then used to draw the phylogenetic relationships among individuals.

## Supplemental Information

Supplemental Information includes Supplemental Data, Supplemental Experimental Procedures, 6 figures and 3 tables.

## Author Contributions

C.U, M.K. A.R.M, and D.B. performed analysis. C.G., M.M., A.R.M, C.U., B.M.H. collected DNA samples. A.B. and S.R. contributed novel analysis tools. P.V.H, M.M, E.H and B.M.H. conceived of the study. C.U., C.R.G, M.M., E.H and B.M.H. wrote the manuscript in collaborations with all co-authors. All authors read and approved of the manuscript.

## Acknowledgements

Funding was provided by a Stanford University CDEHA seed grant to B.M.H. (NIH, NIA P30 AG017253-12) as well as a Stanford University Computation, Evolutionary, and Human Genomics (CEHG) Trainee Research Grant to A.R.M. C.U. was funded by the National Research Foundation of South Africa. C.R.G was funded by T32HG. We thank Jeffrey Kidd for assisting with genotyping of samples. We thank David Poznik for providing off-target mtDNA reads from a separate next-generation sequencing experiment. We thank Aaron Behr and Sohini Ramachandran for pre-publication use of *pong* and Meng Lin for help with analyses. We thank Carlos Bustamante for his encouragement and support of this project. We thank Marcus Feldman for a close reading of our manuscript. We thank Julie Granka, Justin Myrick and Cedric Werely for assistance with saliva sample collection and Ben Viljoen for DNA extractions. Guidance from Ryan Raaum with regards to formulating the surface plots is appreciated. We would like to thank the Working Group of Indigenous Minorities in Southern Africa (WIMSA) and the South African San Institute (SASI) for their encouragement and advice. Finally, we thank Richard Jacobs, Wilhelmina Mondzinger, Hans Padmaker, Willem de Klerk, Hendrik Kaiman, and the communities in which we have sampled; without their support, this study would not have been possible.

